# Effects of Visual Deprivation on Gait Stability During Treadmill Walking: Insights from a Bipedal Model

**DOI:** 10.1101/2025.04.29.651071

**Authors:** Negar Rahimi, Mohsen Amiri, Alireza Kamankesh, Otella Shoja, Lei Zhang, Farnaz Ghassemi, Farzad Towhidkhah

## Abstract

Walking, a fundamental daily activity, relies on sensory input from the somatosensory, visual, and vestibular systems, with vision playing a key role in providing information. When vision is impaired, individuals adjust their walking patterns to maintain stability. This study developed a simplified sagittal model to examine how visual deprivation affects treadmill walking at a controlled speed while ensuring dynamic stability. The model provides a framework for exploring outcomes that are challenging to assess experimentally. We hypothesized that the model would align with experimental data on step frequency, step length, and stability while predicting the steps that could lead to falls without vision. This study presents a bipedal model with two degrees of freedom, controlled by a hierarchical system. Two bio-inspired constraints were implemented: a speed constraint to match with treadmill pace, and a stability constraint to maintain walking stability. The weight of the feedback signal related to joint angles was modified to simulate vision loss. Visual deprivation increased step frequency and decreased step length to prevent falls. Additionally, prolonged deprivation led to step length fluctuations and increased constraint errors, ultimately causing falls. These findings align with human data, validating the potential of the model for rehabilitation research in individuals with visual impairments.

## 1. Introduction

Maintaining stability during human bipedal locomotion requires the coordination of body segments to sustain an intended speed [1-3]. Sensory inputs from the visual, vestibular, and proprioceptive systems play a crucial role in controlling and facilitating normal walking [4]. Although the vestibular, proprioceptive, and sensory feedback provided by muscles contribute to balance [5-7], they are unable to fully compensate for the loss of visual information, leading to an increase in the risk of falls and reducing dynamic stability during walking [4, 8-12]. Accordingly, the visual system plays a primary role in ensuring dynamic stability during locomotion [4, 13-17]. It provides critical information about the surrounding environment, which is essential for detecting and avoiding obstacles [9]. Reducing visual input has been shown to influence key gait parameters, including speed, step length, step width, lower limb kinematics, interlimb coordination, trunk stability, lower-limb symmetry, and activity in the sensorimotor cortex [9, 15, 17-27]. Consequently, there is significant interest in understanding the role of vision in the control of walking, and numerous studies have highlighted its importance in maintaining balance and stability during movement [15, 17, 20, 21, 28-30].

A previous study has included approaches to determine the role of vision in walking overground [31]. However, gait adjustments in the absence of vision differ significantly between treadmill and overground walking, with stride duration being shorter during eyes-closed walking on a treadmill compared with overground walking [26, 32]. In contrast, treadmill walking is crucial since it facilitates postural control in patients with motor disorders by allowing numerous repetitions of rhythmic stepping [26, 32, 33]. Additionally, a treadmill ensures a constant speed during walking and provides a controlled environment for investigating gait features [32, 33]. Nonetheless, none of the previous studies have explicitly modeled and examined the effects of reducing visual input during treadmill walking. To address this, we developed a two-degree-of-freedom bipedal model with a hierarchical control structure to maintain stability in a bipedal model with two key bio-inspired constraints: (1) stability, to ensure dynamic stability was maintained; (2) and speed, so that the model’s pace matched the treadmill speed. These constraints are particularly relevant to treadmill walking, where speed regulation is more critical than in overground walking.

The purpose of this study was to develop a sagittal model to examine the influence of visual deprivation on treadmill walking at a controlled speed while maintaining dynamic stability. The proposed bipedal model employed a hierarchical control structure to evaluate how visual deprivation influenced walking dynamics. Additionally, the model allowed for the exploration of outcomes and parameters that are challenging to measure experimentally. This approach has significant implications for the design of targeted rehabilitation programs for individuals with visual impairments and contributes new knowledge on visual deprivation and the modeling of human movement. We hypothesized that the performance of the model would capture key features of experimental data on how visual deprivation alters walking dynamics, particularly in terms of stability, step frequency, and step length. Additionally, we expected the model to predict the sequence of steps that could lead to a fall in the absence of vision.

## 2. Methods

We proposed a simple model based on a two-degree-of-freedom framework [34-36], which includes both biped dynamics and a hierarchical control system consisting of three-level controllers. It was designed to stabilize walking on a treadmill at a constant speed, preventing the risk of falls caused by fluctuations in walking speed. The controller ensured stability by enforcing the stability constraint and maintaining the model’s center of mass between the two legs. The proposed model was validated using experimental data from participants deprived of visual information while walking on a treadmill.

### 2.1. Hierarchical Control Structure

Figure. 1 shows a hierarchical control structure, comprising three levels: high-, middle-, and low-level controllers. The high-level controller, represented by the green region, defined the desired joint angle trajectory based on constraints involving continuous and discrete dynamics. The input to this control system was the constant speed of the treadmill, with visual feedback further adjusting the joint angles necessary for walking. The middle-level controller coordinated leg displacement by modulating motor commands. Impedance control regulates the interaction between the system and its environment by modeling joint stiffness and damping. This was implemented using a proportional-derivative (PD) controller, which serves as an impedance control by representing the stiffness and damping properties of the joints. The low-level controller, depicted in yellow in Fig. 1, compensates for the nonlinearity of the dynamic by feedback linearization. A feedback loop simulating visual deprivation, based on previous studies, [26, 28, 29, 37] impacts both movement strategy and controller coefficients.

**Fig. 1.**
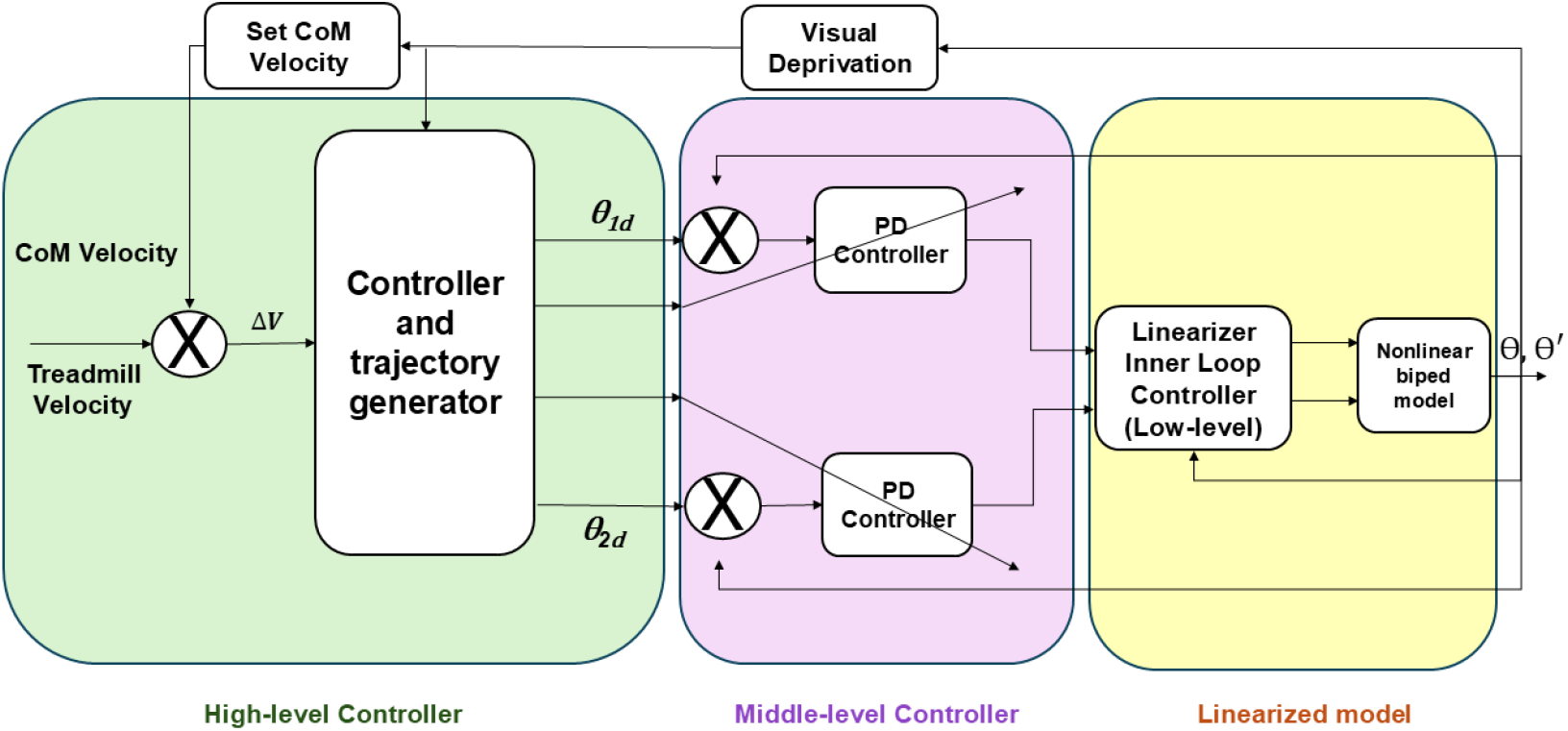
Diagram of the model architecture for bipedal locomotion during visual deprivation during treadmill walking. CoM denotes the center of mass, and the PD controller refers to the proportional-derivative controller. The angles *θ*_*1d*_ and *θ*_*2d*_ represent the desired angle of the support leg relative to the vertical axis and the desired angle between the two legs, respectively. ΔV denotes the difference between the treadmill velocity and the CoM velocity.

### 2.2. Experimental protocol and setup

Ten individuals (four males and six females; age: 31.5 ± 3.8 years) participated in the experiment. Each participant walked along the middle of a motorized treadmill at a preferred speed (1.09 ± 0.18 m/s) without handrails while wearing a safety harness. All individuals had normal or corrected-to-normal vision with no history of neurological, musculoskeletal, or orthopedic conditions. To simulate the absence of vision, participants wore liquid crystal glasses that turned opaque or transparent at the toe-off phase of the right leg. Gait steps were analyzed during three phases: before deprivation (vision intact), during deprivation (vision blocked), and after deprivation (vision restored). Spatiotemporal gait variables were recorded with a three-dimensional motion-capture system (Optotrak Certus, Northern Digital Inc, Canada) that tracked a heel marker. Eight steps from each visual phase were analyzed for each participant.

### 2.3. Bipedal Walker Model

A two-dimensional (2D) bipedal walker model was used, focusing on sagittal stability and step length (Fig. 2). The model included two legs with two degrees of freedom (DOF), assuming zero hip width with concentrated mass at the end of each leg. To simulate walking dynamics, one leg consistently maintained contact with the treadmill surface. Both discrete and continuous equations were incorporated, augmented by collision equations to capture the impact dynamics to enhance the overall model accuracy.

**Fig. 2.**
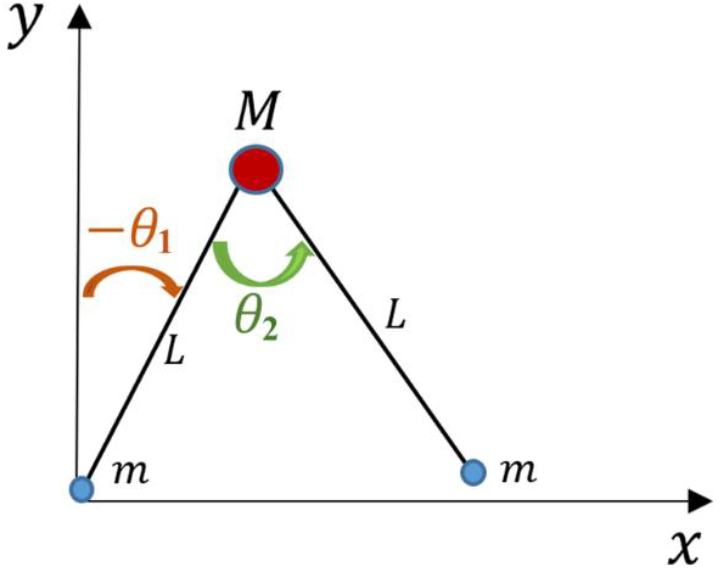
Diagram of the bipedal model. *θ*_*1*_ represents the angle of the support leg relative to the vertical axis, whereas *θ*_*2*_ indicates the angle between the two legs. *M* represents the concentrated mass of the upper body. Each leg was depicted as a link with length *L* and mass *m* at its distal end.

#### 2.3.1. Dynamic and Control

Hybrid dynamics were utilized by integrating both discrete and continuous equations, augmented by collision equations to capture the impact dynamics to enhance the overall model accuracy [38].

The continuous dynamics of the system were governed by Lagrange’s equation [39], as shown in Eq. 1:

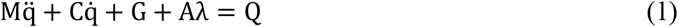

where *M* represented the inertia/mass matrix, *C* denoted the matrix of Coriolis or centrifugal terms, and *G* was the vector of gravitational forces and torques. *A* defined the constraint associated with the foot compliance, whereas *λ* served as the Lagrange multiplier that provided the forces needed to maintain the foot constraint. Additionally, Q corresponded to the vector of generalized forces and torques acting on the system.

In this work, the foot was assumed to remain fixed at the point of contact, preventing sliding during foot strike, with the collision modeled as inelastic. Lagrange’s method was used to calculate the velocity immediately after each foot strike, incorporating discrete collision equations. While foot angles before and after contact remained unchanged, the velocity differed between steps, causing different speeds for the primary and subsequent steps. These equations were derived from the energy differences between before and after collision states, with the heel speed reaching zero at foot strike, stabilizing it through restraining forces (Eq. 2) [36]:

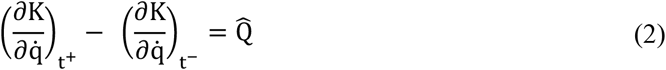

where *K* is the kinetic energy, *q* is the configuration vector consisting of the joint angles, and 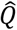 is the generalized impulse due to non-conservative forces, such as damping. Throughout each step, the PD coefficients remained constant. This simplification reduced model complexity; however, these coefficients should ideally vary continuously.

The control approach was developed with notable features, including its ability to compensate for variations in step length under visual deprivation, its time-invariant nature, and its capacity to adjust both step length and frequency. The equations (Eq. 3 to 5) illustrate how the controller enforced the speed and stability constraints through the parameter “*h*”:

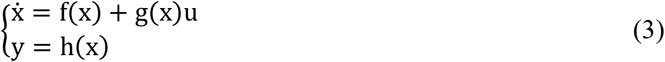

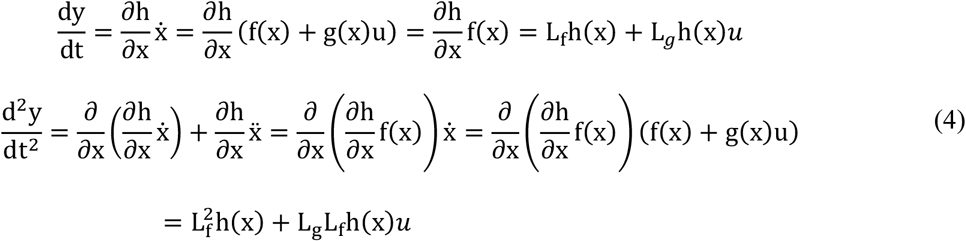

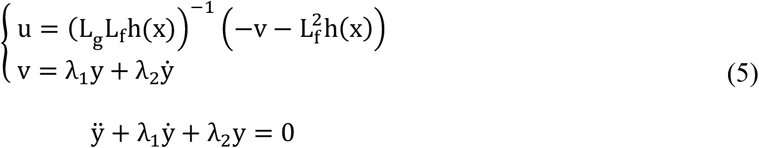

where *x* represents the state vector of the system, *f*(*x*) describes the system dynamics without control input, while *g*(*x*) shows how the control input *u* affects the system. *u* is the control input, and *y* is the system’s output, with *h*(*x*) as the output function. An auxiliary input *v* is introduced as a virtual control variable to impose the desired output behavior. The parameters *λ*_*1*_ *and λ*_*2*_ are positive design constants that serve as the coefficients of the first derivative and the output terms, respectively, in the closed-loop dynamics. These parameters are selected to ensure system stability and to regulate the transient response characteristics, including damping and convergence speed.

The constraints represented by *L*_*f*_*h* and *L*_*g*_*h* (Eq. 4) are Lie derivatives of the output function *h*(*x*) along the system dynamics *f*(*x*) and control distribution *g*(*x*) respectively. These terms describe how the output evolves under the system’s natural dynamics and control inputs. The speed and stability constraints served two primary functions: (1) maintaining stability by ensuring the center of mass remained consistently positioned between the two legs, and (2) synchronizing the speed of the model to treadmill speed. Without these constraints, the model’s stability would not be ensured, even during simple steps with normal visual input. The following equations define the stability constraint on the hip joint (Eq. 6):

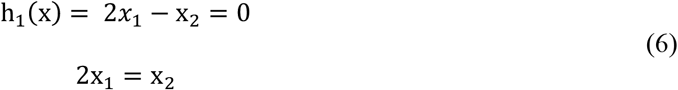

Where *x*_1_ and *x*_2_ represent the joint angles *θ*_1_ and *θ*_2_, where *θ*_1_ corresponds to the ankle angle and *θ*_2_ corresponds to the hip angle. The stability condition defined the required location of the CoM to ensure the model maintained stability. Finally, the constraints on the ankle joint were expressed through equations corresponding to specified walking speeds (Eq. 7):

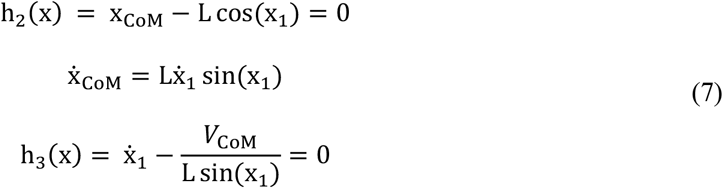

Exceeding the treadmill speed could lead to a forward fall, whereas lagging treadmill speed resulted in a backward fall. To prevent these outcomes, this equation (Eq. 7) guaranteed that the speed of the model matched the treadmill speed (*V*_*CoM*_). By incorporating these constraints, the model generated a trajectory with appropriate introduced angles. To maintain the constraint within the correct range, two coefficients (*λ*_1_and *λ*_2_in Eq. 5) were introduced for additional adjustment. Additionally, two errors velocity and stability were continuously monitored.

#### 2.3.2. Visual Deprivation

To examine the influence of vision, the weight of the feedback signal has been changed in the model (Fig.1). Since this feedback directly impacts the constraints (as shown in Eq. 6 and 7), the influence of visual deprivation on walking was investigated by examining the change in model performance. Using a proportional gain concept, two scenarios were established: the first scenario, with normal vision, maintained a full feedback gain of 1.0 for both proprioception and vision; the second scenario, simulating the absence of vision, reduced the feedback gain to 0.6.

## 3. Results

### 3.1. Parameter Estimation

The simulations were conducted using MathWorks MATLAB Simulink (Version R2024a). The parameters used in the model are presented in Table 1. Although several parameters, including treadmill speed, upper body mass, concentrated mass of each leg, and leg length, were adapted from those reported in a previous study [40, 41], some parameters were adjusted to meet the specific requirements of our study (Table 1).

**Table 1.** Model parameters used in the simulations.

| Parameter | Description | Value | Unit |
| --- | --- | --- | --- |
| M | Upper body mass | 31.73 | kg |
| m | Concentrated mass of each leg | 13.5 | kg |
| $I_1, I_2$ | Inertia momentum | 0.2 | kg.m<sup>2</sup> |
| G | Gravitational acceleration | 9.81 | m/s<sup>2</sup> |
| $V_d$ | Treadmill constant speed | 0.2 | m/s |
| $L_1, L_2$ | Leg length | 1 | m |
| $K_{ps}$ | Ankle proportional gain | 220 | N. m/rad |
| $K_{pt}$ | Hip proportional gain | 220 | N. m/rad |
| $K_{ds}$ | Ankle derivative gain | 42 | N. m/rad |
| $K_{dt}$ | Hip derivative gain | 14 | N. m. s/rad |

Within the controller, the coefficients *K*_*ds*_ and *K*_*ps*_ were associated with ankle control, and *K*_*dt*_ and *K*_*pt*_ were related to hip control.

### 3.2. The influence of vision on gait characteristics

Table 2 reports the mean and standard deviation for step length, step frequency, swing time, and relative speed across three conditions (before, during, and after visual deprivation) for both experimental and simulation results. To ensure comparability with experimental data, the modeling values were scaled by a constant factor [42] of 3.35. Visual deprivation reduced step length and increased step frequency compared with the conditions before and after deprivation. Additionally, swing times and relative speed were greater before and after deprivation than during visual deprivation, closely aligning with the experimental findings (Table 2).

**Table 2.** Mean and standard deviation of step length, step frequency, swing time, and relative speed before, during, and after deprivation conditions in experimental test and proposed model.

| Test | Parameters | Before visual<br>deprivation | During visual<br>deprivation | After visual<br>deprivation |
| --- | --- | --- | --- | --- |
| Experiment | Step Length (cm) | $59.89 \pm 7.48$ | $57.3 \pm 7.49$ | $59.4 \pm 6.79$ |
| | Step Frequency (step/min) | $55.80 \pm 4.14$ | $64.61 \pm 4.63$ | $55.59 \pm 6.43$ |
| | Swing Time (s) | $0.38 \pm 0.02$ | $0.37 \pm 0.02$ | $0.39 \pm 0.02$ |
| | Relative Speed (m/s) | $1.00 \pm 6 \times 10^{-3}$ | $0.99 \pm 0.02$ | $1.01 \pm 0.02$ |
| Model | Step Length (cm) | 58.48 | 58.43 | 58.47 |
|  | Step Frequency (step/min) | 55.03 | 55.11 | 54.99 |
|  | Swing Time (s) | 0.3897 | 0.3890 | 0.3899 |
|  | Relative Speed (m/s) | 1.086 | 1.084 | 1.09 |

### 3.3. Kinematic Tracking: Trajectory and Errors

Figure 3 illustrates the joint-angle trajectory, CoM velocity, and the speed and stability errors during a single step. The angle trajectories of *θ*_*1*_ and *θ*_*2*_ are depicted (Fig. 3 A). The consistent ratio of these angles indicated that the CoM remained stable between the two legs, thereby ensuring dynamic stability. Additionally, when the initial CoM speed exceeded the treadmill speed limit, the model promptly adjusted its velocity to 0.2 m/s to match the pace of the treadmill (Fig. 3 B). These adjustments reflected the ability of the model to control movement and adhere to the imposed constraints. Constraint errors during treadmill walking represent velocity and stability errors (Fig. 3 C). The figure depicts that while errors initially increased as the model began walking, the control system effectively reduced them to zero (Fig. 3 C). This observation confirmed the proper functioning of the stability constraint and ensured the maintenance of stability throughout the treadmill movement. Additionally, it validated the effective operation of the speed constraint, demonstrating that the model remained stable on the treadmill despite fluctuations in speed.

**Fig. 3.**
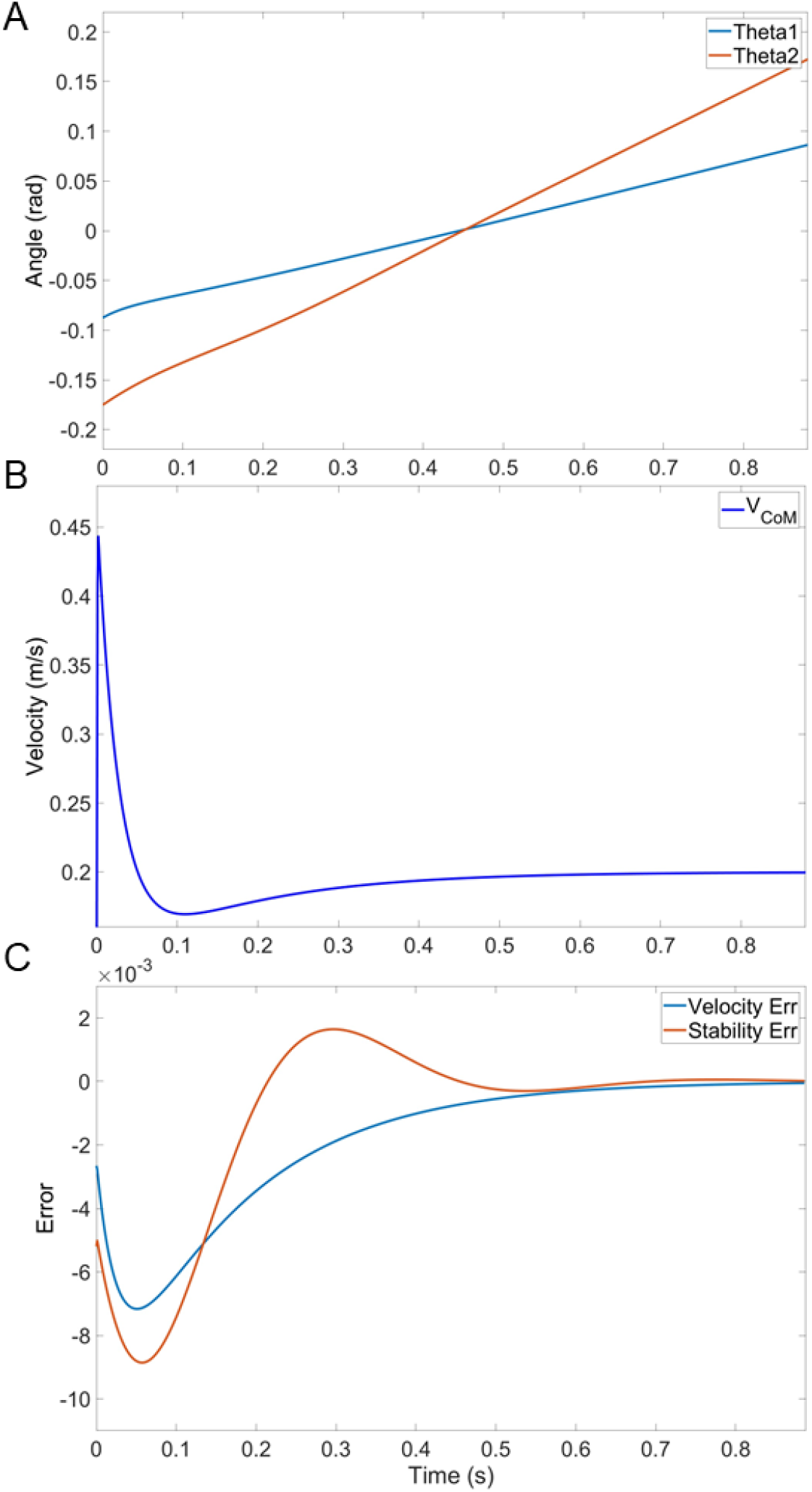
Trajectories and errors during a single step on the treadmill without visual deprivation. A) Angle trajectories of *θ*_*1*_ (blue) and *θ*_*2*_ (orange); B) velocity of the center of mass (*V*_*CoM*_); and C) Errors (Err) for speed and stability constraint, shown in blue and orange, respectively.

### 3.4. Influence of vision and control parameters on step length

Figure. 4 shows the model step length before, during, and after visual deprivation while walking on the treadmill. Prior to visual deprivation, the model maintained relatively constant step length across steps (Fig. 4). Visual deprivation was introduced at the 8th step and lasted for approximately 4 steps, initially causing a slight alteration in step length. Subsequent adjustments led to a further decrease in step length. Vision was restored at step 16, prompting the model to compensate and gradually return to the before-deprivation step length over the following 4 steps (Fig. 4).

**Fig. 4.**
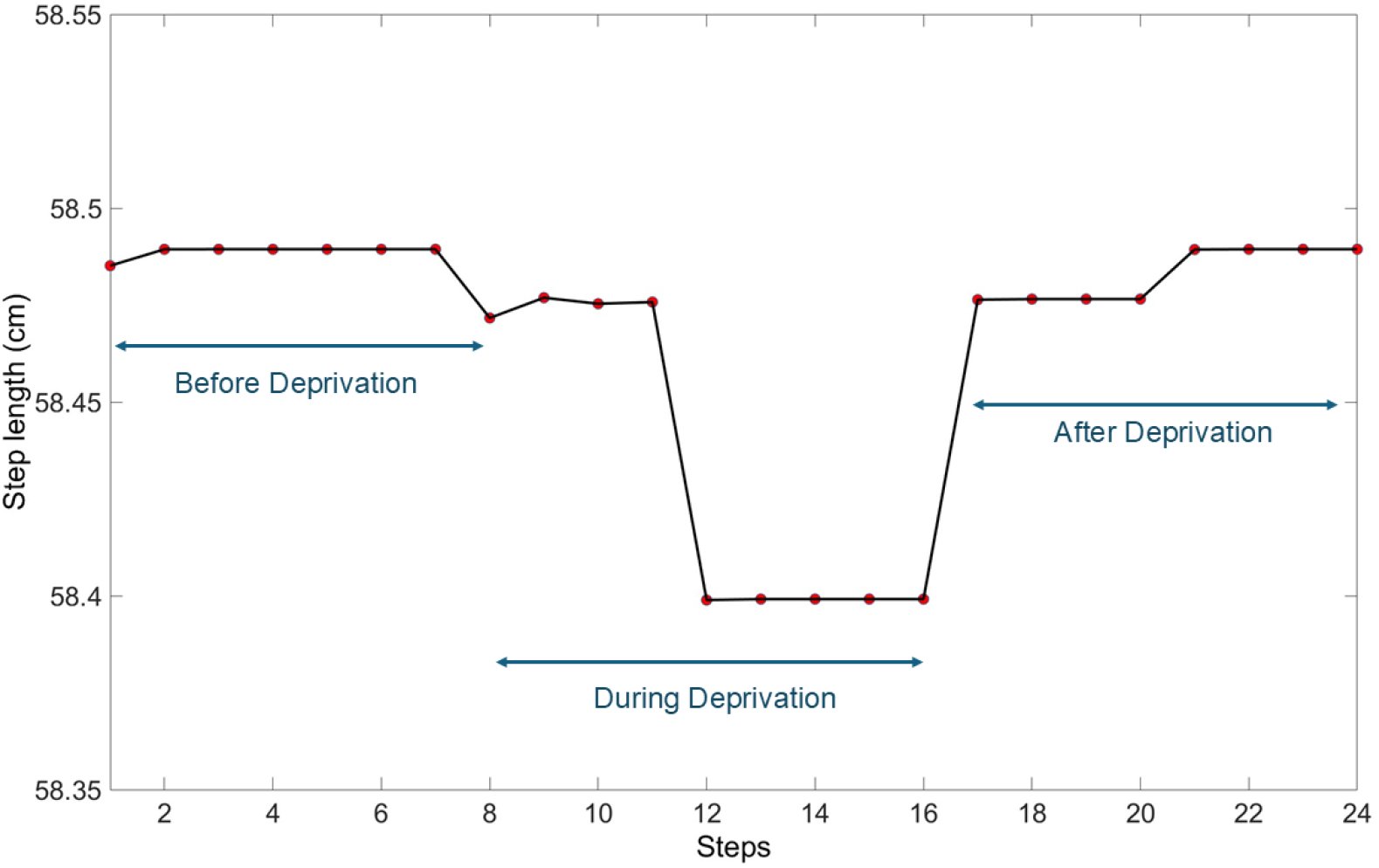
Model step length before, during, and after visual deprivation. Each red dot represents an individual step.

Furthermore, the effect of prolonged visual deprivation on treadmill walker indicated that after approximately 18 steps, the angular trajectory began to deviate, and the CoM velocity exhibited significant alterations. Consequently, velocity and stability errors failed to revert to their initial states. The accumulation of these errors compromised dynamic stability, ultimately causing a loss of stability and a subsequent fall.

## 4. Discussion

This paper introduces a stable model of bipedal treadmill walking under visual deprivation. Although several studies have investigated the impact of visual deprivation on stability during treadmill walking [20, 28-30, 43], relatively few have focused on modeling this phenomenon. Visual deprivation disrupts higher-level processing, leading to incomplete information from proprioception and other sensory feedback systems [26, 28, 29, 37]. Because proprioceptive and other sensory systems cannot fully compensate for the absence of visual input, individuals are compelled to adopt an alternative walking strategy [4, 6, 8], which we examined by developing a hierarchical model to vary the level of visual feedback.

We implemented two key constraints: stability and speed. Dingwell et al. [44] emphasized that both speed and position control are crucial for meeting the fundamental requirement of staying on the treadmill. The stability constraint ensured walking stability by maintaining the center of mass between the legs, regulated through the ankle and hip joint angles [45]. Additionally, the speed constraint maintained a constant speed to prevent forward or backward displacement, ensuring the individual would not fall off the treadmill. As Dingwell et al. further suggest, participants must maintain the same average position in the middle of the treadmill and match walking speed to the constant pace of the treadmill to avoid falling [44], a point also emphasized in previous studies on speed consistency for safety and stability [32, 33]. Typically, individuals select a preferred walking speed to optimize energy expenditure and ensure efficient locomotion [33]. However, the constant speed of the treadmill may alter walking strategies and constrain users to a straighter path than overground walking [32]. Gait speed, a fundamental aspect of walking, acts as an adaptive mechanism that can respond to perturbations and enhance dynamic stability. Therefore, abrupt changes in gait speed can disturb gait symmetry, leading to unstable joint movement patterns and compromised stability.

The results indicated that the model maintained a constant step length and walking speed while adhering to both speed and stability constraints during full-vision condition. Step length remained relatively unchanged for the initial steps in the absence of vision, which is consistent with the conclusion of Leech et al. that the human nervous system quickly recalls walking patterns to navigate changing environments [46]. However, the subsequent changes in step length were evident in our experimental data and are consistent with previous studies [10, 47, 48]. Our experimental findings indicated that decreases in step length during visual deprivation resulted in a walking speed that was less than the pace of the treadmill. In response, step frequency increased, which aligns with the results of Espy et al. [48] of individuals adopting shorter and faster steps to ensure stability in the absence of vision. Individuals attempted to return to a normal cadence within the first few steps after reopening their eyes and regaining awareness of their position on the treadmill. Maintaining a consistent position on the treadmill requires matching the average walking speed to the treadmill’s pace [44], which promotes stability and prevents falls by reducing step length and increasing step frequency. Within a few subsequent steps following deprivation, participants returned to a stable state, achieving the same step length as before deprivation and matching treadmill speed. These findings are also supported by previous studies [12, 29, 49, 50], which reported temporary reductions in step length during visual deprivation, followed by recovery once visual input was restored.

The model adhered to speed and stability constraints, with their errors converging to zero within a brief period before and after visual deprivation. Substantial fluctuations in stability and speed errors arose due to step-to-step variations in step length, persisting across multiple consecutive steps [44]. However, error correction and individual effort were compromised during visual deprivation. Destabilizing and blindfolding experiments reported in the literature [9, 49] similarly show that compensatory mechanisms fail under such conditions, leading to falls. After prolonged visual deprivation, the model fell off the treadmill due to accumulating errors, speed discrepancies relative to the treadmill, and an inability to maintain dynamic stability. In contrast, during the experimental study, the test was stopped at the first signs of instability to ensure participant safety.

### 4.1 Limitations and Future Directions

Several limitations in our study could be acknowledged and addressed in future work. First, the proposed model utilizes a two degrees of freedom structure with a simplified design. While this design omits certain anatomical features, such as knee movement, it was chosen deliberately to reduce computational complexity and focus on key walking dynamics. These simplifications allow for a more focused analysis of visual deprivation effects, while still a reasonable level of accuracy in simulating human performance. Secondly, the current study did not investigate the double support phase, as the model was designed for an initial investigation of visual deprivation effects on walking dynamics. Future studies could also include this phase in their model. Finally, the coefficients in the model were updated discretely, once per step. However, adjusting these coefficients continuously would be beneficial to have more realistic models.

## 5. Conclusion

We demonstrated a two-degree-of-freedom bipedal model with a hierarchical control system capable of replicating stable walking on a treadmill under visual deprivation. The model integrated two bio-inspired constraints designed to maintain dynamic stability while ensuring the model’s speed is synchronized with the treadmill’s velocity. Simulation results show that visual deprivation induced adjustments in the walking strategy, characterized by a reduction in step length and an increase in step frequency. Additionally, the model predicted that approximately 18 steps of visual deprivation could lead to a fall. These findings can inform studies on the effects of visual deprivation on treadmill walking, the design of robotic systems, and future work in modeling and rehabilitating individuals with visual impairments.

## References

[1] H.B. Menz, S.R. Lord, R.C. Fitzpatrick, Acceleration patterns of the head and pelvis when walking on level and irregular surfaces, Gait Posture 18(1) (2003) 35–46. DOI 10.1016/S0966-6362(02)00159-5

[2] S. van Andel, A.R. Schmidt, P.A. Federolf, Distinct coordination patterns integrate exploratory head movements with whole-body movement patterns during walking, Sci. Rep. 13(1) (2023) 1235. DOI 10.1038/s41598-022-26848-x

[3] S.M. Bruijn, P. Meyns, I. Jonkers, D. Kaat, J. Duysens, Control of angular momentum during walking in children with cerebral palsy, Res. Dev. Disabil. 32(6) (2011) 2860–2866. DOI 10.1016/j.ridd.2011.05.019

[4] M.S. Redfern, L. Yardley, A.M. Bronstein, Visual influences on balance, J. Anxiety Disord. 15(1-2) (2001) 81–94. DOI 10.1016/S0887-6185(00)00043-8

[5] A. Kamankesh, N. Rahimi, I.G. Amiridis, C. Sahinis, V. Hatzitaki, R.M. Enoka, Distinguishing the Activity of Flexor Digitorum Brevis and Soleus Across Standing Postures with Deep Learning Models, Gait Posture (2025). DOI 10.1016/j.gaitpost.2024.12.014

[6] S.-Y. Cho, Y.-U. Ryu, H.D. Je, J.H. Jeong, S.-Y. Ma, H.-D. Kim, Effects of illumination on toe clearance and gait parameters of older adults when stepping over an obstacle: a pilot study, J. Phys. Ther. Sci. 25(3) (2013) 229–232. DOI 10.1589/jpts.24.229

[7] A.E. Patla, T.C. Davies, E. Niechwiej, Obstacle avoidance during locomotion using haptic information in normally sighted humans, Exp. Brain Res. 155 (2004) 173–185. DOI 10.1007/s00221-003-1714-z

[8] A. Goodworth, K. Perrone, M. Pillsbury, M. Yargeau, Effects of visual focus and gait speed on walking balance in the frontal plane, Hum. Mov. Sci. 42 (2015) 15–26. DOI 10.1016/j.humov.2015.04.004

[9] M. Iosa, A. Fusco, G. Morone, S. Paolucci, Effects of visual deprivation on gait dynamic stability, Sci. World J. 2012(1) (2012) 974560. DOI 10.1100/2012/974560

[10] A. Hallemans, E. Ortibus, F. Meire, P. Aerts, Low vision affects dynamic stability of gait, Gait Posture 32(4) (2010) 547–551. DOI 10.1016/j.gaitpost.2010.07.018

[11] R.F. Reynolds, B.L. Day, Visual guidance of the human foot during a step, J. Physiol. 569(2) (2005) 677–684. DOI 10.1113/jphysiol.2005.095869

[12] F. Reynard, P. Terrier, Role of visual input in the control of dynamic balance: variability and instability of gait in treadmill walking while blindfolded, Exp. Brain Res. 233 (2015) 1031–1040. DOI 10.1007/s00221-014-4177-5

[13] A.E. Patla, How is human gait controlled by vision, Ecol. Psychol. 10(3-4) (1998) 287–302. DOI 10.1080/10407413.1998.9652686

[14] D.S. Marigold, Role of peripheral visual cues in online visual guidance of locomotion, Exerc. Sport Sci. Rev. 36(3) (2008) 145–151. DOI 10.1097/jes.0b013e31817bff72

[15] O. Shoja, F. Towhidkhah, H. Hassanlouei, M.F. Levin, A. Bahramian, S. Nadeau, et al., Reaction of human walking to transient block of vision: analysis in the context of indirect, referent control of motor actions, Exp. Brain Res. 241(5) (2023) 1353–1365. DOI 10.1007/s00221-023-06593-x

[16] S. Rietdyk, C.K. Rhea, Control of adaptive locomotion: effect of visual obstruction and visual cues in the environment, Exp. Brain Res. 169 (2006) 272–278. DOI 10.1007/s00221-005-0345-y

[17] O. Shoja, M. Shojaei, H. Hassanlouei, F. Towhidkhah, M. Amiri, H. Boroomand, et al., Lack of visual information alters lower limb motor coordination to control center of mass trajectory during walking, J. Biomech. 155 (2023) 111650. DOI 10.1016/j.jbiomech.2023.111650

[18] A. Hallemans, S. Beccu, K. Van Loock, E. Ortibus, S. Truijen, P. Aerts, Visual deprivation leads to gait adaptations that are age-and context-specific: I. Step-time parameters, Gait Posture 30(1) (2009) 55–59. DOI 10.1016/j.gaitpost.2009.02.018

[19] R. Moe-Nilssen, J.L. Helbostad, T. Akra, L. Birkedal, H.A. Nygaard, Modulation of gait during visual adaptation to dark, J. Motor Behav. 38(2) (2006) 118–125. DOI 10.3200/JMBR.38.2.118-125

[20] O. Shoja, A. Farsi, F. Towhidkhah, A.G. Feldman, B. Abdoli, A. Bahramian, Visual deprivation is met with active changes in ground reaction forces to minimize worsening balance and stability during walking, Exp. Brain Res. 238 (2020) 369–379. DOI 10.1007/s00221-020-05722-0

[21] O. Shoja, M. Shojaei, H. Hassanlouei, F. Towhidkhah, L. Zhang, Quantifying Human Gait Symmetry During Blindfolded Treadmill Walking, Motor Control. 1(aop) (2024) 1–16. DOI 10.1123/mc.2023-0028

[22] B. Kadirvelu, C. Gavriel, S. Nageshwaran, J.P.K. Chan, S. Nethisinghe, S. Athanasopoulos, et al., A wearable motion capture suit and machine learning predict disease progression in Friedreich?-≥s ataxia, Nat. Med. 29(1) (2023) 86–94. DOI 10.1038/s41591-022-02159-6

[23] M.S.B. Hossain, Z. Guo, H. Choi, Estimation of lower extremity joint moments and 3d ground reaction forces using imu sensors in multiple walking conditions: A deep learning approach, IEEE J. Biomed. Health Inform. 27(6) (2023) 2829–2840. DOI 10.1109/JBHI.2023.3262164

[24] J.F. O’Brien, R.E. Bodenheimer, G.J. Brostow, J.K. Hodgins, Automatic joint parameter estimation from magnetic motion capture data, arXiv preprint arXiv:2303.10532 (2023). DOI 10.20380/GI2000.09

[25] K. John, J. Stenum, C.-C. Chiang, M.A. French, C. Kim, J. Manor, et al., Accuracy of video-based gait analysis using pose estimation during treadmill walking versus overground walking in persons after stroke, Phys. Ther. 104(2) (2024) pzad121. DOI 10.1093/ptj/pzad121

[26] A.S. Oliveira, B.R. Schlink, W.D. Hairston, P. Konig, D.P. Ferris, Restricted vision increases sensorimotor cortex involvement in human walking, J. Neurophysiol. 118(4) (2017) 1943–1951. DOI 10.1152/jn.00926.2016

[27] N. Rahimi, A. Kamankesh, I.G. Amiridis, S. Daneshgar, C. Sahinis, V. Hatzitaki, R.M. Enoka, Distinguishing among standing postures with machine learning-based classification algorithms, Exp. Brain Res. 243(1) (2025) 1–13. DOI 10.1007/s00221-024-06959-9

[28] M.J. Dunn, S.K. Rushton, Lateral visual occlusion does not change walking trajectories, J. Vis. 18(9) (2018) 11–11. DOI 10.1167/18.9.11

[29] F. Saucedo, F. Yang, Effects of visual deprivation on stability among young and older adults during treadmill walking, Gait Posture 54 (2017) 106–111. DOI 10.1016/j.gaitpost.2017.03.001

[30] C. Tobar, E. Martinez, N. Rhouni, S.-J. Kim, The effects of visual feedback distortion with unilateral leg loading on gait symmetry, Ann. Biomed. Eng. 46 (2018) 324–333. DOI 10.1007/s10439-017-1954-x

[31] J. Ondřej, J. Pettré, A.-H. Olivier, S. Donikian, A synthetic-vision based steering approach for crowd simulation, ACM Transactions on Graphics (TOG) 29(4) (2010) 1–9. DOI 10.1145/1778765.1778860

[32] J.H. Hollman, M.K. Watkins, A.C. Imhoff, C.E. Braun, K.A. Akervik, D.K. Ness, A comparison of variability in spatiotemporal gait parameters between treadmill and overground walking conditions, Gait Posture 43 (2016) 204–209. DOI 10.1016/j.gaitpost.2015.09.024

[33] C. Park, K. Park, Dynamic Stability of Human Walking in Response to Sudden Speed Changes, Symmetry. 16(1) (2023) 26. DOI 10.3390/sym16010026

[34] M. Garcia, A. Chatterjee, A. Ruina, M. Coleman, The simplest walking model: stability, complexity, and scaling, (1998). DOI 10.1115/1.2798313

[35] J.M. Morawski, A simple model of step control in bipedal locomotion, IEEE Trans. Biomed. Eng. (6) (1978) 544–549. DOI 10.1109/TBME.1978.326289

[36] T. McGeer, Dynamics and control of bipedal locomotion, J. Theor. Biol. 163(3) (1993) 277–314. DOI 10.1006/jtbi.1993.1121

[37] D.P. et al, Book Neuroscience, John Wiley 2018.

[38] W.-L. Ma, A.D. Ames, From bipedal walking to quadrupedal locomotion: Full-body dynamics decomposition for rapid gait generation, 2020 IEEE International Conference on Robotics and Automation (ICRA), IEEE, 2020, pp. 4491–4497.

[39] J. Ginsberg, Engineering dynamics, Cambridge University Press 2008.

[40] R.D. Gregg, J.W. Sensinger, Towards biomimetic virtual constraint control of a powered prosthetic leg, IEEE Trans. Control Syst. Technol. 22(1) (2013) 246–254. DOI 10.1109/TCST.2012.2236840

[41] S. Plagenhoef, F.G. Evans, T. Abdelnour, Anatomical data for analyzing human motion, Res. Q. Exerc. Sport. 54(2) (1983) 169–178. DOI 10.1080/02701367.1983.10605290

[42] M. Garcia, A. Chatterjee, A. Ruina, Efficiency, speed, and scaling of two-dimensional passive-dynamic walking, Dynamics and Stability of Systems 15(2) (2000) 75–99. DOI 10.1080/713603737

[43] J. Hao, T.W. Buster, G.M. Cesar, J.M. Burnfield, Virtual reality augments effectiveness of treadmill walking training in patients with walking and balance impairments: a systematic review and meta-analysis of randomized controlled trials, Clin. Rehabil. 37(5) (2023) 603–619. DOI 10.1177/02692155221138309

[44] J.B. Dingwell, J.P. Cusumano, Identifying stride-to-stride control strategies in human treadmill walking, PLoS One. 10(4) (2015) e0124879. DOI 10.1371/journal.pone.0124879

[45] L. Tesio, V. Rota, The motion of body center of mass during walking: a review oriented to clinical applications, Front. Neurol. 10 (2019) 999. DOI 10.3389/fneur.2019.00999

[46] K.A. Leech, R.T. Roemmich, A.J. Bastian, Creating flexible motor memories in human walking, Sci. Rep. 8(1) (2018) 94. DOI 10.1038/s41598-017-18538-w

[47] E. D’Hondt, V. Segers, B. Deforche, S.P. Shultz, A. Tanghe, I. Gentier, et al., The role of vision in obese and normal-weight children’s gait control, Gait Posture 33(2) (2011) 179–184. DOI 10.1016/j.gaitpost.2010.10.090

[48] D.D. Espy, F. Yang, T. Bhatt, Y.-C. Pai, Independent influence of gait speed and step length on stability and fall risk, Gait Posture 32(3) (2010) 378–382. DOI 10.1016/j.gaitpost.2010.06.013

[49] P.M. McAndrew, J.M. Wilken, J.B. Dingwell, Dynamic stability of human walking in visually and mechanically destabilizing environments, J. Biomech. 44(4) (2011) 644–649. DOI 10.1016/j.jbiomech.2010.11.007

[50] S.M. O’Connor, A.D. Kuo, Direction-dependent control of balance during walking and standing, J. Neurophysiol. 102(3) (2009) 1411–1419. DOI 10.1152/jn.00131.2009

